# Defining and detecting global transcriptional amplitude in circadian gene expression

**DOI:** 10.1101/2025.11.14.688559

**Authors:** Michael Saint-Antoine, Ron C. Anafi

## Abstract

Many genes exhibit circadian rhythms in expression. The amplitude of oscillation, both in core clock and circadian output genes, may differ from person to person. Mutations in core clock genes are known to alter global rhythmic properties, and researchers often informally discuss “circadian amplitude.” Yet, it remains unclear whether, in the general population, differences in transcriptional amplitude are largely gene-specific, or if they reflect a global, transcriptome-wide pattern -- whether some individuals have globally higher or lower amplitude across the set of all rhythmic genes.

We used Cosinor regression to reanalyze four human skin time-series transcriptomic datasets (paired epidermis/dermis samples, N=11, N=19) and found that, using either absolute or relative amplitude measures, distributions of gene amplitudes tended to cluster by subject. Using a non-parametric, permutation-based statistical test, we found that in many subjects this global amplitude trend was statistically significant (*p* ≤ 0.01). Furthermore, we found that when rhythmic genes were divided into two sets based on peak time (genes peaking before-noon and after-noon), the subjects’ global amplitude in one set predicted global amplitude in the other set (*p* ≤ 0.05). We also found that in the paired epidermis/dermis datasets, subjects’ global amplitude in epidermis predicted their global amplitude in the dermis (*p* ≤ 0.05).

After identifying these trends in the skin datasets, we then found that evidence for subject-specific transcriptional rhythm strength replicated across 6 additional human time-course datasets from adipose, muscle, and blood. Perhaps surprisingly, across datasets, we found that neither established metrics of core clock transcriptional organization nor the amplitude of core clock transcription were strongly correlated with subject-specific global amplitude.

## Introduction

Many aspects of mammalian physiology exhibit circadian (almost daily) rhythms, including easily observed changes in the sleep-wake cycle, body temperature, feeding behavior, and attention (1). Less outwardly visible changes in cellular and molecular physiology are similarly common. Specific metabolic or immune functions within an organ or tissue may vary with the 24-hour day/night cycle. At the cellular level, these circadian rhythms are controlled by a molecular clock involving a transcriptional/translational feedback loop (2). These autonomous, self-sustaining cellular clocks are also modulated by a central clock in the suprachiasmatic nucleus (SCN) of the hypothalamus, helping to maintain inter-cellular and inter-tissue coherence (3,4). Although the core molecular clock mechanism involves relatively few genes, circadian rhythms can also be detected and quantified in the expression of many of their downstream targets.

Modern techniques such as RNA-seq and microarray experiments allow researchers to study whole-transcriptome circadian variation. For example, Zhang et al. report results from a multi-tissue mouse model, finding that 43% of all protein-coding genes showed circadian oscillations in at least one tissue (5), and Littleton et al. analyzed tissue-level differences in the fraction of rhythmic transcripts and overall amplitude of the circadian transcriptome (6). Ruben et al. found that, across 13 human tissues, ∼7,000 genes showed circadian expression in at least one tissue, including many that encode drug targets, transporters, or metabolizing enzymes (7). Indeed, recent work has focused on the influence of circadian time on the efficacy and toxicity of immune checkpoint inhibitors (8,9), and mechanistic data show that time-of-day-dependent CD8+ T-cell tumor infiltration modulates immune checkpoint inhibitor efficacy (10). Work from our own lab has suggested that altered molecular rhythms may have both functional and prognostic implications in Luminal A breast cancer (11) and Alzheimer’s disease (12).

Given the evidence supporting the therapeutic relevance of circadian rhythms, there is increasing interest in how these rhythms vary between individuals and thus may influence the personalization of circadian medicine. Indeed, recent studies have quantified inter-individual differences in the expression of core clock genes in human skin (13), although quantifying inter-individual differences in amplitude across the entire rhythmic transcriptome has received less attention. The potential for global descriptors of circadian variation is of both clinical and basic scientific interest. We specifically questioned if there is a global, subject-specific strength in circadian transcriptional rhythms within the general, healthy population. Simply put, if a person has relatively high amplitude in some circadian genes compared to other people, do they tend to also have relatively high amplitude in other genes? Does it make sense to say that someone has high-amplitude or low-amplitude rhythms in gene expression *in general?* We sought to address this question using publicly available transcriptomic time-series datasets involving different people.

## Methods

### Datasets

Our primary analysis utilized four datasets involving time-course collections of human skin samples:

The first two skin datasets we reanalyzed were epidermis and dermis time-series microarray datasets (GEO ID GSE205155) from Del Olmo et al. (14), hereafter referred to as EPIDERMIS_1 and DERMIS_1. Eleven healthy young adults (6 male, 5 female, age 20-30) were sampled via 3mm punch biopsies from the back every 4 hours over 24 hours (7 time points per subject). Epidermis and dermis were separated and gene expression profiled by microarray.

We similarly analyzed the epidermis (EPIDERMIS_2, GEO ID GSE139301) and dermis (DERMIS_2, GEO ID GSE139300) time-series microarray datasets reported by Wu et al. (15,16). These datasets describe gene expression from 2mm forearm punch biopsies of healthy male subjects, age 21-49. After excluding one subject with only 3 available samples, this left us with data from 19 subjects, each with 4 biopsy samples each taken at 6 hour intervals. Epidermis and dermis were isolated by laser capture microdissection, and gene expression was profiled by microarray.

Subsequent analysis included data from other tissues, including white adipose (referred to as ADIPOSE) reported by Christou et al. (17) with GEO ID GSE87761, and skeletal muscle (referred to as MUSCLE) reported by Perrin et al. (18) with GEO ID GSE108539. In addition, we evaluated several whole blood datasets: Braun et al. (19) with GEO ID GSE113883, Arnardottir et al. (20) with GEO ID GSE56931, referred to as BLOOD_1 and BLOOD_2, respectively, and the whole-blood dataset of Möller-Levet et al. (21). Möller-Levet and colleagues evaluated the same participants both after completing 7 nights of sufficient sleep opportunity (10 hours time-in-bed/night) and 7 nights of comparatively insufficient sleep (6 hours time-in-bed/night). Both conditions were immediately followed by a 39-41 hour constant-routine protocol during which subjects stayed awake and samples were collected. In our analysis, we used only the baseline data from the first 24 hours of samples for each subject. We refer to the measurements following the week of sufficient sleep as BLOOD_3_NORMAL, and the measurements following the week of insufficient sleep as BLOOD_3_SR (“SR” for “sleep-restriction”). Across all of these studies, participants were recruited as generally healthy adults without reported sleep disorders or shift-work schedules. **Table 1** shows a brief summary of these datasets.

**Table 1:**
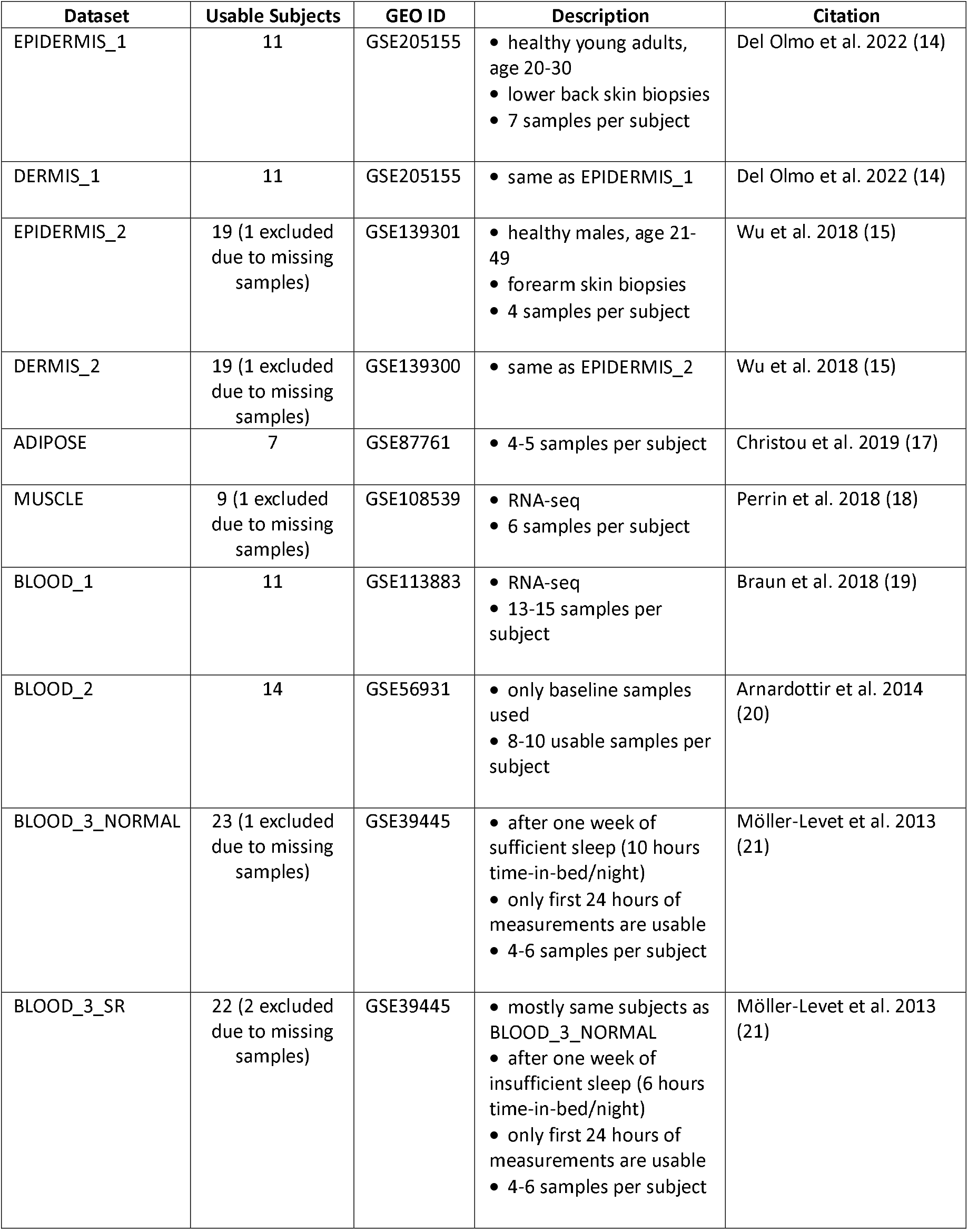
Brief summary of information for all 10 datasets.

### Data processing

While we attempted to utilize a common analysis framework, inherent differences between the datasets required data-set specific processing steps. For example, MUSCLE and BLOOD_1 are RNA-seq datasets, and required different processing from the others, which are microarray datasets.

We first removed any subjects with fewer than 4 samples. Next, we removed probes with mean expression < 4. Probe level data was then unlogged and probe-set measurements were mapped to the corresponding genes using platform specific mappings in GEO. In cases where multiple probes mapped to the same gene, we took their mean. In cases where a probe did not map to an annotated gene symbol, we kept the probe ID as the identifier and still included it in the processed data. The top 10,000 genes based on mean expression were retained for further analysis (excepting EPIDERMIS_2 which had <10,000 transcripts after processing).

The processing pipeline resulted in standardized inputs for our downstream Cosinor analysis: a linear-scale, non-negative gene expression matrix with rows corresponding to the top ∼10,000 most highly-expressed genes and columns for each sample. Metadata with the subject identifier and timepoint of collection for each sample were stored in separate arrays. R scripts used for data processing and subsequent analyses are available at https://github.com/ranafi/Subject-Global-Amplitude.

### Cosinor analysis

We used Cosinor regression (22,23) to detect and model rhythms in gene expression for each dataset in a two-step approach. We first modeled rhythms across all subjects collectively to identify transcripts rhythmic at the population-level (called the “collective Cosinor analysis”). Then for each transcript assessed as cycling at the population we refit a Cosinor model to each subject individually to estimate subject-specific amplitude and mean expression (called the “subject-specific Cosinor analysis”).

For the collective Cosinor analysis, time-course transcript expression *x* (*t*) was fit to two nested linear models for each gene:

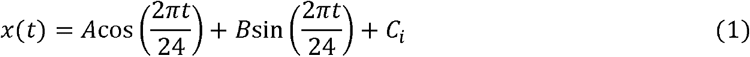

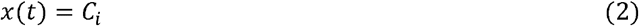

with *i* ∈ { 1,2, …, *N* } denoting one of the *N* subjects and *C*_*i*_ representing the subject-specific circadian-corrected mean, called a MESOR (midline estimating statistic of rhythm) in this context. The parameters *A* and *B* were fit across all subjects collectively. The fit of the nested models was compared with an F-test, applied using the *anova()* function in R, yielding a p-value for each gene. P-values were then adjusted for multiple testing using the method of Benjamini and Hochberg (24), and are referred to as q-values after adjustment. We then identified significantly rhythmic (*q* ≤ 0.05) genes for further analysis. For these rhythmic genes, we also calculated population-level absolute amplitude, relative amplitude, and acrophase using the fit Cosinor parameters:

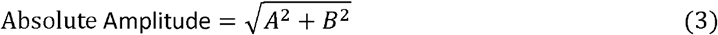

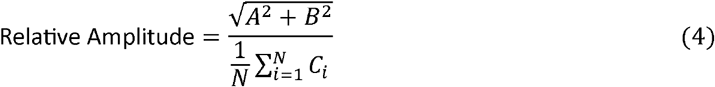

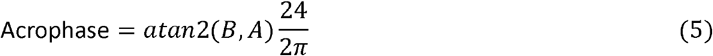

Based on these population-level rhythmic properties, we classified genes as “high-amplitude” if they had a relative amplitude ≥ 0.1, and also recorded their population-level acrophase for use in a subsequent analysis. **Table 2** shows a summary of the collective Cosinor results for each dataset.

**Table 2:** Cosinor analysis results.

| Dataset | Total Genes | Rhythmic Genes | High-Amplitude Genes | Genes Peaking 0-12h | Genes Peaking 12-24h | Rhythmic Core Cock Genes |
| --- | --- | --- | --- | --- | --- | --- |
| EPIDERMIS_1 | 10,000 | 3,783 | 2,789 | 1852 | 1931 | ARNTL, CRY1, CRY2, DBP, NFIL3, NPAS2, NR1D2, PER1, PER2, PER3 |
| DERMIS_1 | 10,000 | 4,520 | 2,206 | 2419 | 2101 | ARNTL, BHLHE41, CRY1, DBP, NFIL3, NR1D1, NR1D2, PER1, PER2, PER3, TEF |
| EPIDERMIS_2 | 9,849 | 3,194 | 981 | 1609 | 1585 | ARNTL, BHLHE41, CLOCK, CRY1, CRY2, DBP, HLF, NFIL3, NPAS2, NR1D2, PER1, PER2, PER3, RORA, RORC, TEF |
| DERMIS_2 | 10,000 | 202 | 158 | 119 | 83 | ARNTL, BHLHE41, CLOCK, HLF, NFIL3, NPAS2, NR1D2, PER1, PER2, PER3, TEF |
| ADIPOSE | 10,000 | 593 | 519 | 292 | 301 | ARNTL, BHLHE41, CRY2, HLF, NFIL3, NR1D1, NR1D2, PER1, PER2 |
| MUSCLE | 10,000 | 2,435 | 2,151 | 1431 | 1004 | BHLHE41, CIART, CLOCK, CRY2, DBP, HLF, NFIL3, NR1D1, NR1D2, PER1, PER2, PER3, RORA, RORC, TEF |
| BLOOD_1 | 10,000 | 7,132 | 4,755 | 5113 | 2019 | ARNTL, CRY2, NR1D1, PER1, TEF |
| BLOOD_2 | 10,000 | 4,246 | 426 | 2397 | 1849 | ARNTL, DBP, NFIL3, NR1D1, NR1D2, PER2, PER3, RORA |
| BLOOD_3_NORMAL | 10,000 | 4,091 | 1,108 | 2066 | 2025 | DBP, NFIL3 |
| BLOOD_3_SR | 10,000 | 2,332 | 468 | 1272 | 1060 | CRY2, NFIL3 |

Next, we performed the subject-specific Cosinor analysis, this time fitting the Cosinor model to each rhythmic gene for each subject individually, allowing us to estimate a subject-specific amplitude for each gene. We calculated the subject-specific relative amplitude as the absolute amplitude divided by the MESOR for each subject. Then, for each gene we performed a rank-normalization, transforming the subject-specific relative amplitudes to percentile scores with:

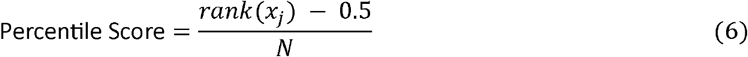

Where *rank(x*_*j*_) is the rank order of the relative amplitude for the *j*-th subject for the given gene. The result of this pipeline was that for each dataset with *m* rhythmic genes and *n* subjects, we recorded an *m* by *n* matrix:

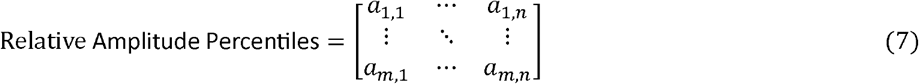

where each element *a*_*i*,*j*_ represents the relative amplitude percentile score for the *i*-th gene, in the *j*-th subject. In other words, this matrix tells us how high or low of a relative amplitude each subject had for a gene, compared to the other subjects. For convenience, we will refer to these relative amplitude percentile scores as simply “amplitude” throughout this paper, and this matrix (called the “amplitude matrix”) is the primary object of analysis in the Results section of this paper. Amplitude matrices, as well as matrices containing the non-normalized absolute and relative amplitudes for each subject, were saved as RDS objects for further analysis.

## Results

### Transcriptome-wide distributions of amplitudes differed visually between subjects, with robust differences in means

For a given dataset, each column of the amplitude matrix contains the distribution of amplitudes for a particular subject across all rhythmic genes. We sought to determine if these distributions differ from subject to subject in a meaningful way. **Figure 1A** shows histogram plots of the amplitude distributions, for the four skin datasets, with subjects ordered according to means of the distributions, from highest to lowest. Each histogram shows the number of transcripts for each subject at the specified percentile rank. **Figure 1B** shows bar plots of the means of these amplitude percentile ranks for each subject. We note the striking visual differences in both the distributions and means, especially when comparing the subjects at the high and low extremes.

**Figure 1:**
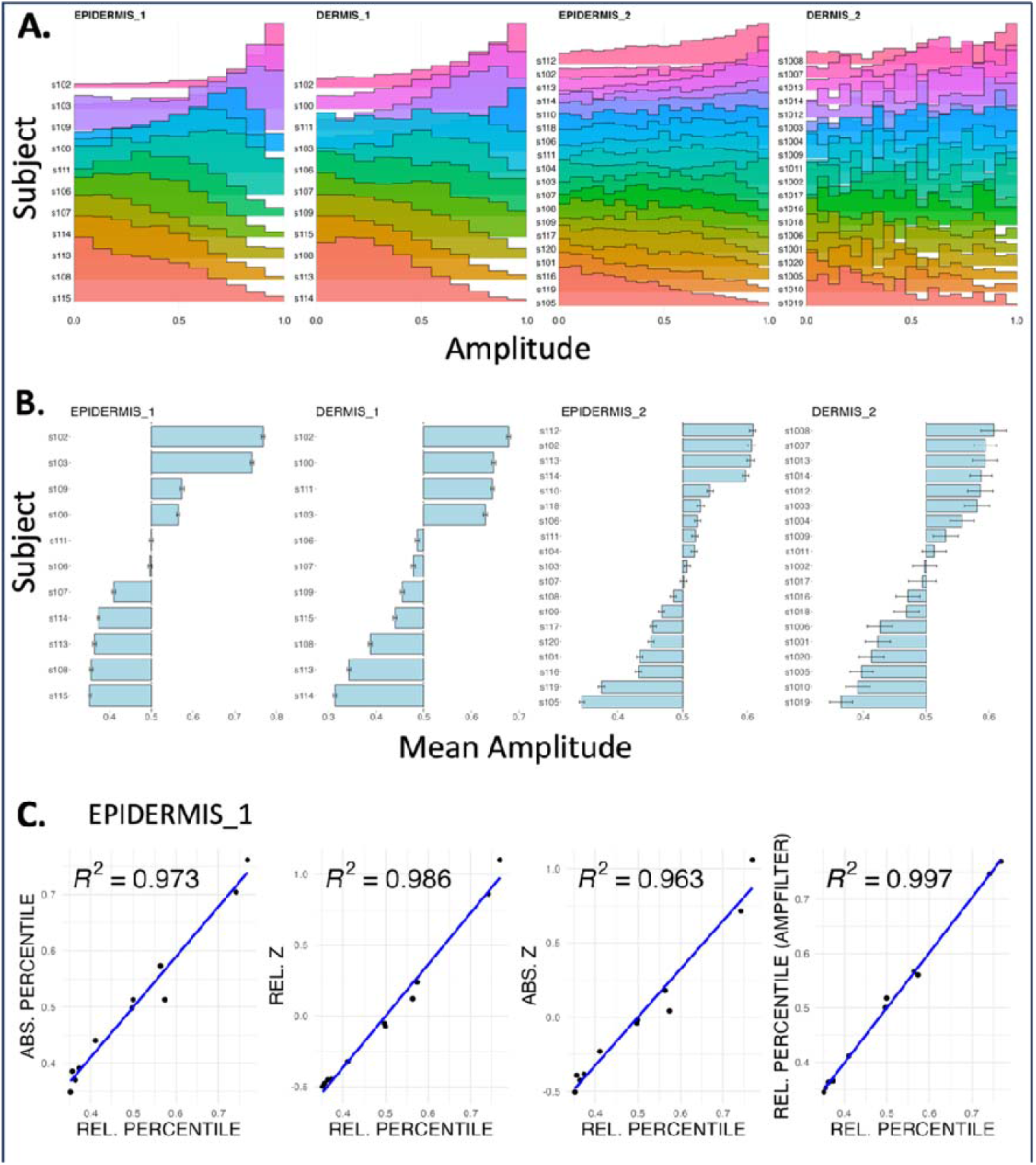
Subject specific trends in transcript amplitude across four skin datasets. **A**.) We used Cosinor regression to estimate the subject-specific relative amplitude for each rhythmic gene. These plots show the distributions of percentile-normalized relative amplitudes (referred to as simply “amplitude” for convenience) for each subject, in each of the four skin datasets, with subjects ordered by their mean amplitude from highest to lowest. The differences between the subject amplitude distributions are visually apparent, especially when comparing subjects at opposite extremes (for example, in EPIDERMIS_1 subject s115’s amplitude distributions skews lower, while subject s102’s distribution skews higher). **B**.) The means of the subject-specific amplitude distributions shown in **Fig. 1A** are plotted, with bars showing standard error, for the four skin datasets. Again, we note the visible differences between subjects, especially those at opposite extremes. **C**.) Subject rankings are robust to a variety of amplitude metrics: We repeated the analysis using relative (rather than absolute) amplitudes from the Cosinor regression, percentile (rather than z-score) normalization, and calculating the subject-specific amplitude of all of the rhythmic genes, rather than considering only high-amplitude genes. The correlation between subject averaged metrics are plotted. Subject-specific amplitudes metrics are very closely regardless of metric.

As noted in the Methods section, we calculated subject-specific amplitudes using a percentile normalization of relative amplitudes for each gene, considering all rhythmic genes. Importantly, subject-specific amplitude means were robust to different judgement calls made in this calculation (**Figure 1C**): z-score (rather than percentile) normalization, using absolute (rather than relative) amplitudes, and using only high-amplitude genes (rather than all rhythmic genes). From this point on, we will use the term “global amplitude” to refer to a subject’s mean amplitude across all rhythmic genes, calculated using percentile-normalized relative amplitudes from the subject-specific Cosinor analysis. However, for several of the analyses presented in this paper, the Supplementary Information includes results using z-score normalization, absolute amplitudes, and considering high-amplitude genes only.

### Many subjects showed a statistically significant trend in global amplitude

Due to the stochastic nature of gene expression, variation in measured amplitude between subjects is expected for any given rhythmic gene, as well as for the average amplitude across all rhythmic genes. The key question of this study is whether the variation in transcriptional amplitude between subjects across the set of all rhythmic genes is entirely random, or if there is an underlying trend to it, such that some subjects tend to have higher or lower amplitude in general across the rhythmic transcriptome relative to the other subjects.

We used permutation-based statistics to formally test this. Consider again the amplitude matrix (Equation 7), which has a row for every gene and a column for every subject, with each element *a*_*i*,*j*_ representing the amplitude for the *i* -th gene, in the *j* -th subject. The null hypothesis is that the rows (genes) in this matrix are independent of each other. In other words, if we know that Subject A has a higher amplitude than Subject B in Gene 1, that does not give us any information about which subject has a higher amplitude in Gene 2. The goal of the statistical test is to identify the subject with the most extreme global amplitude (either higher or lower compared to the other subjects), and compute how likely we would be to observe a global amplitude that extreme under the null hypothesis.

The workflow for the statistical test is as follows. We begin with the amplitude matrix for a particular dataset, and calculate the column means, giving us the global amplitude for each subject. We then select the subject whose global amplitude is the furthest from the 50^th^ percentile (computed as |*x* − 0.5| where *x* is the subject’s global amplitude percentile average). To evaluate the significance of this metric, we use bootstrapping to generate a null distribution for comparison. We randomly shuffle each row of the amplitude matrix independently, so that for each gene the amplitudes are randomly assigned to the subjects in a way that is independent of the other genes. Again, we calculate the column means of this matrix to generate a simulation of global amplitudes under the null hypothesis and select the most extreme in terms of distance from the mean (|*x* − 0.5| for each global amplitude *x*). We save this value as a measure of global amplitude extremeness under the null hypothesis. We repeat this process many times (1000 total), each time reshuffling each row of the amplitude matrix, to generate a test distribution under the null hypothesis.

Then, we compare our observed value to our test distribution. If the observed value falls in the top 1% of the test distribution, we consider that subject to have a statistically significant global amplitude trend (*p* ≤ 0.01), meaning that their global amplitude is either higher or lower than 0.5 to an extent that would be unlikely to be observed due to chance alone under the null hypothesis. We then delete that subject’s column from the amplitude matrix and recompute the percentiles for each gene using the amplitudes of the remaining subjects. This ensures that each subsequent test evaluates extremeness relative to an updated cohort that is no longer influenced by previously identified outliers. For example, if all subjects were similar except for one individual who had the highest amplitude for every transcript, including that subject in the reference distribution would cause all other subjects to appear artificially low in their average ranks. After removing the most extreme subject, we repeat the test for the next most extreme individual. This process continues until the most extreme remaining subject is no longer statistically different from the updated cohort, or until only one subject remains. **Figure 2** illustrates this workflow.

**Figure 2:**
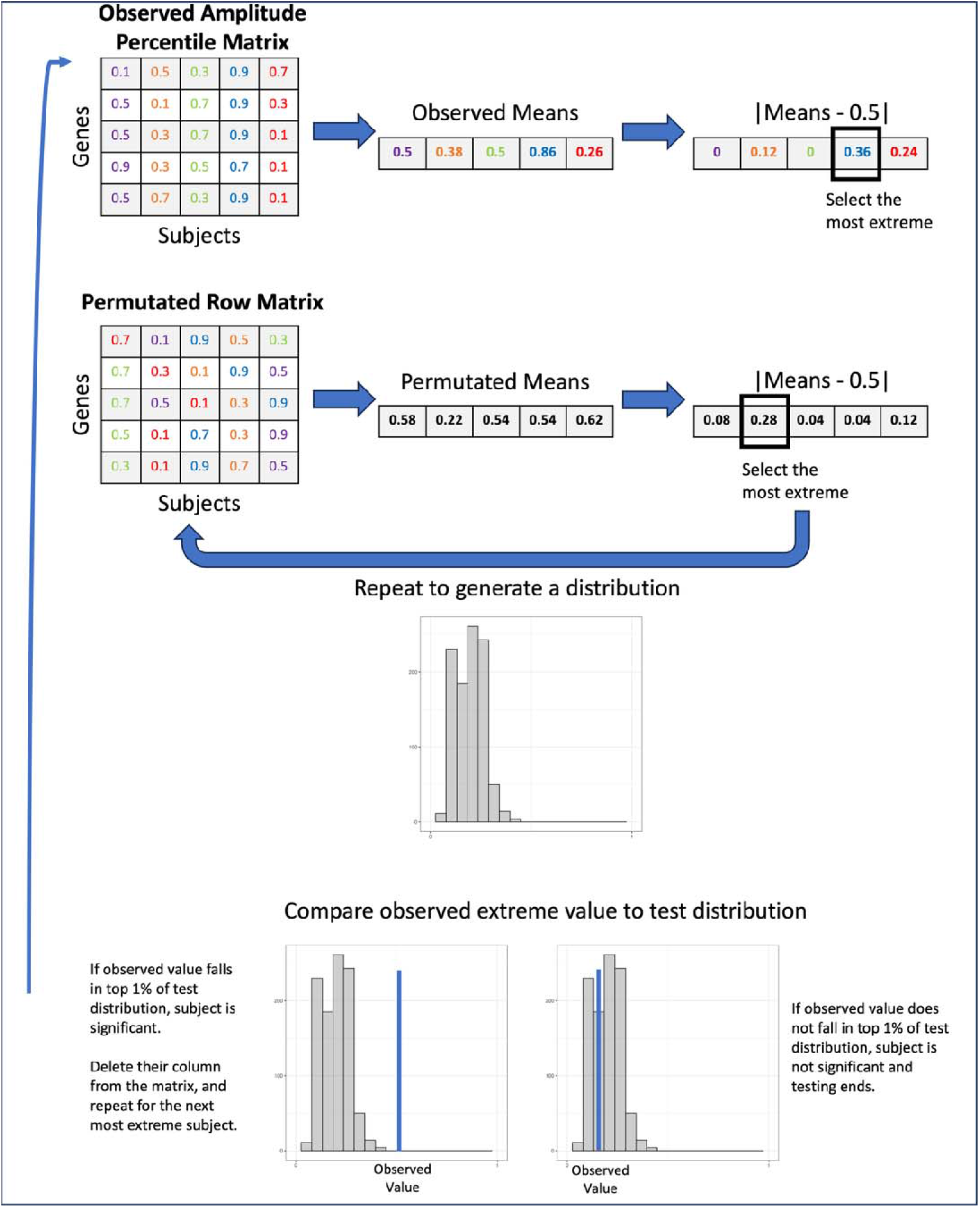
Diagram of permutation test workflow. We begin with the amplitude matrix, and calculate the mean for each subject. We then identify the subject whose mean is the further from 0.5, and call this distance the *observed value* (computed as where is the subject’s mean amplitude). Then, we randomly shuffle each row of the amplitude matrix independently of each other, recalculate the column means, and compute the distance of the most extreme mean from 0.5. We repeat this 1000 times total to generate a test distribution. We then compare the observed value to the test distribution. If it falls in the top 1%, then we consider that subject to be statistically significant (*p* ≤ 0.01), delete their column of data from the amplitude matrix, and repeat the process of the second most extreme subject (renormalizing the rows for every iteration of the test). We repeat this process until a subject is identified as non-significant, at which point the test ends, or until there is only one subject remaining, who is then considered significant by default.

**Table 3** shows how many subjects were identified as having a statistically significant trend in global amplitude, for each of the four skin datasets. Indeed, in three of the datasets the majority of subjects were found to have significant differences in global amplitude, while in the remaining dataset (DERMIS_2) 8 out of 19 subjects did. The Supplementary Information section of this paper also includes results using z-score (rather than percentile) normalization, absolute (rather than relative) amplitudes, and considering only high amplitude genes (rather than all rhythmic genes). We found that that the overall results (both in terms of the relative rankings of subjects and number of subjects found to have statistically different global amplitude metrics when compared to the broader cohort) were generally consistent regardless of the specific metric used.

**Table 3:**
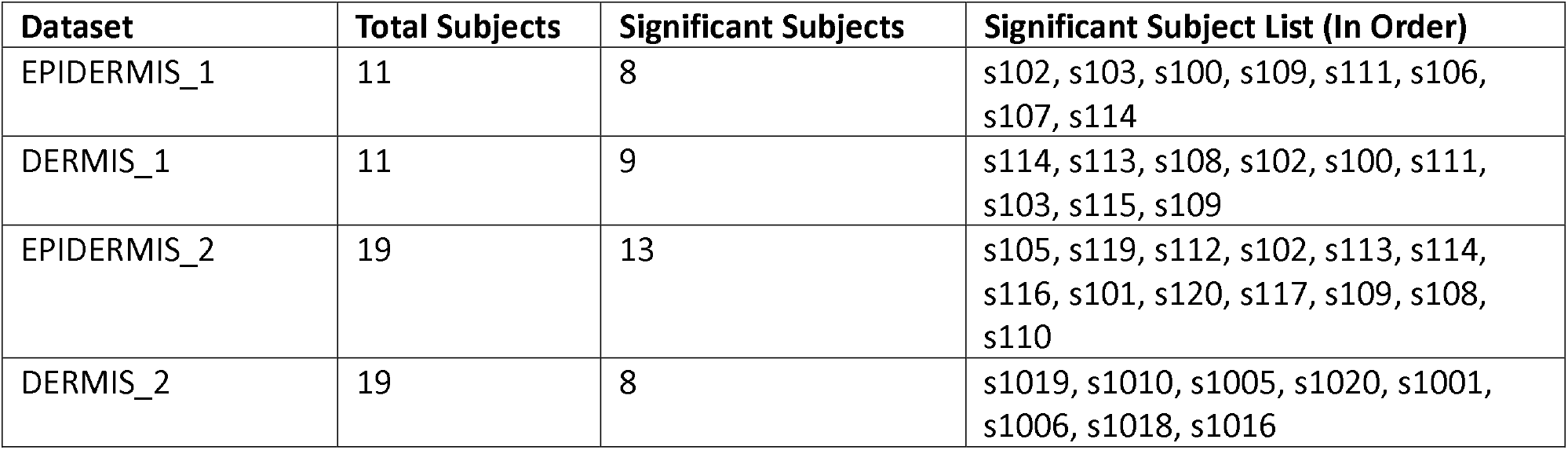
Permutation test results for the four skin datasets.

### Subject-specific global amplitude was correlated between before-noon-peaking and after-noon-peaking genes, and between epidermis and dermis

Rhythmic genes showed bimodal acrophase distributions in 3/4 skin datasets (**Figure 3A**), with the only exception being DERMIS_2, which had fewer total rhythmic genes than the others. We split the genes by acrophase into two groups: those with before-noon peak time (0 - 12h), and those with after-noon peak time (12 - 24h), based on the population-level phase estimates from the collective Cosinor analysis. We then independently computed the subject-specific global amplitudes for both before-noon and after-noon gene sets. Across subjects, before-noon and after-noon global amplitude were strongly correlated (**Figure 3B**) in EPIDERMIS_1 (*R*^*2*^ = 0.847, *p* = 5.883304e-05), DERMIS_1 (*R*^*2*^ = 0.855, *p* = 4.566727e-05), and EPIDERMIS_2 (*R*^*2*^ = 0.843, *p* = 2.947722e-08). In DERMIS_2, the correlation was weaker but still statistically significant (*R*^*2*^ = 0.426, *p* = 0.002).

**Figure 3:**
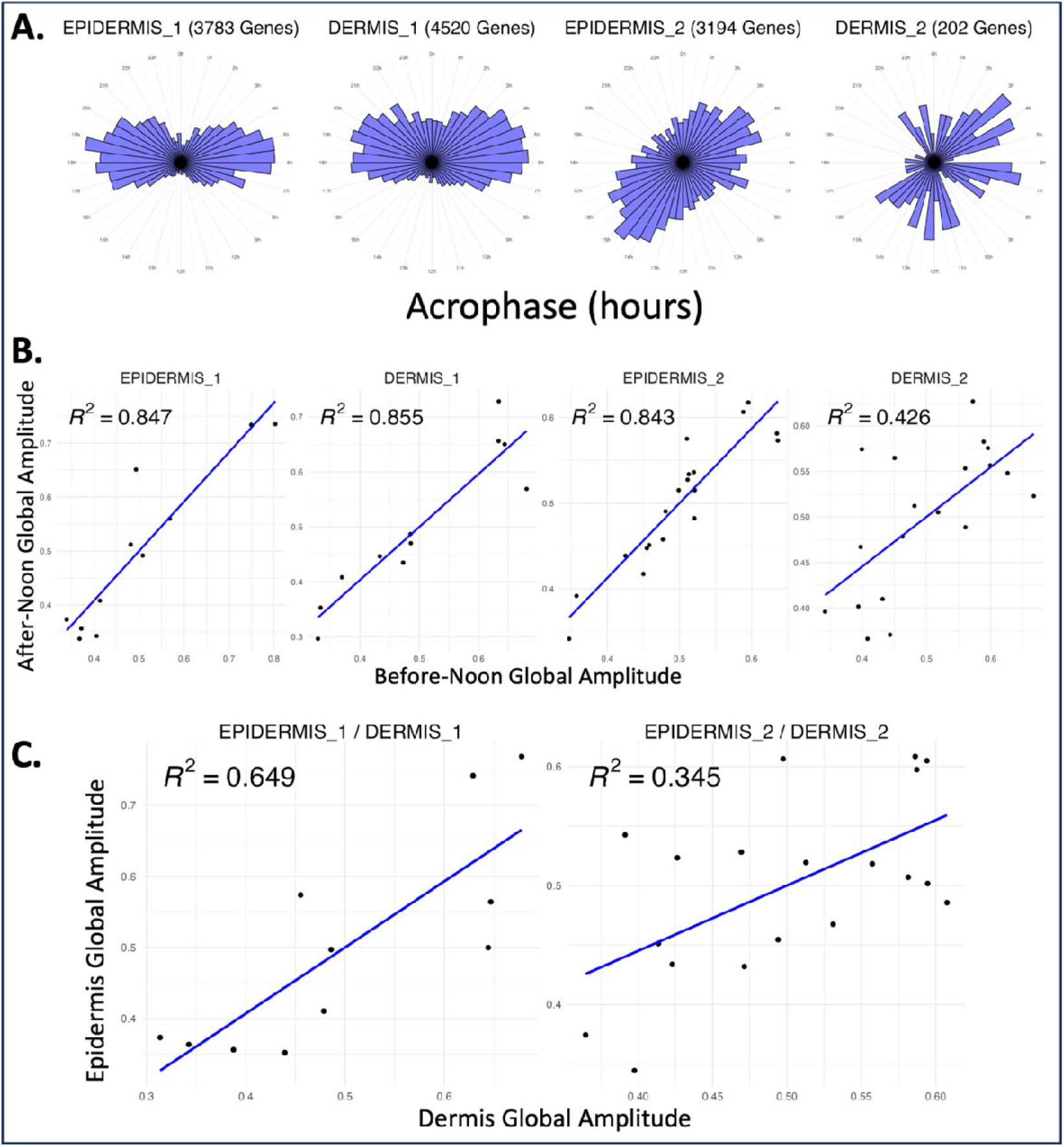
Correlation of subject specific global transcriptional amplitude using distinct transcript sets or cell-types. **A**.) Rose plots depict the distribution of acrophases (computed in the collective Cosinor analysis, across all subjects) for the rhythmic genes in each dataset. Three of the four skin datasets show bimodal acrophase distributions, with the only exception being DERMIS_2, which had fewer rhythmic genes than the others. **B**.) For each dataset, we split the rhythmic genes into two groups: those with before-noon acrophases (0 - 12h), and those with after-noon acrophases (12 - 24h). We then computed the subject-specific mean amplitudes in these two groups separately, to see if subject’s mean amplitude for genes peaking before-noon predicted their mean amplitude for genes peaking after-noon. These correlations were quite strong in EPIDERMIS_1, DERMIS_1, and EPIDERMIS_2, and weaker but still statistically significant (*p* = 0.002) in DERMIS_2. **C**.) The DERMIS_1/EPIDERMIS_1 datasets contain dermis and epidermis samples for the same group of subjects, as do DERMIS_2/EPIDERMIS_2. These plots show that the subjects’ mean amplitude in the dermis predicts their mean amplitude in the epidermis, in both DERMIS_1/EPIDERMIS_1 (*p* = 0.002) and DERMIS_2/EPIDERMIS_2 (*p* = 0.008).

We also evaluated the correlation between subject-specific global amplitude, comparing measures from epidermis and dermis for the same subjects (**Figure 3C**). We found that subject-specific global amplitude was well correlated between the two skin layers (*R*^*2*^ = 0.649, *p* = 0.002 in EPIDERMIS_1/DERMIS_1; *R*^*2*^ = 0.345, *p* = 0.008 in EPIDERMIS_2/DERMIS_2), suggesting that subjects with higher epidermal rhythm strength also had higher dermal rhythm strength.

### These results replicated in six additional datasets from adipose, muscle, and blood

To see if these patterns were evident in non-skin tissues, we replicated our analysis in the adipose (17), muscle (18), and blood (19–21) datasets (**Table 4**). As with the skin datasets, we note that most of the subjects in each dataset had a statistically significant difference in global transcript amplitude as compared to the collection of tested subjects. **Table 5** shows the correlation between the global subject amplitude rankings when using exclusively either the genes with before-noon or after-noon peak times. In each dataset (excepting BLOOD_1) the two rankings are well correlated. However, we also found that in the BLOOD_3_NORMAL and BLOOD_3_SR datasets, subjects’ global amplitude in the “NORMAL” condition (sufficient sleep) did *not* predict their global amplitude in the “SR” condition (restricted sleep) (*R*^*2*^ = 0.019, *p* = 0.58), suggesting that subject-specific global amplitude trends may be responsive to behavioral changes.

**Table 4:** Permutation test results for the six additional datasets.

| Dataset | Total Subjects | Significant Subjects |
| --- | --- | --- |
| ADIPOSE | 7 | 4 |
| MUSCLE | 9 | 6 |
| BLOOD_1 | 11 | 11 |
| BLOOD_2 | 14 | 12 |
| BLOOD_3_NORMAL | 23 | 20 |
| BLOOD_3_SR | 22 | 18 |

**Table 5:** Morning-peak amplitude vs evening-peak amplitude for the six additional datasets.

| Dataset | $R^2$ | $p$ | Correlation Direction |
| --- | --- | --- | --- |
| ADIPOSE | 0.715 | 0.0165 | Positive |
| MUSCLE | 0.982 | 2.497071e-07 | Positive |
| BLOOD_1 | 0.135 | 0.2672 | Positive |
| BLOOD_2 | 0.893 | 3.597969e-07 | Positive |
| BLOOD_3_NORMAL | 0.764 | 5.09451e-08 | Positive |
| BLOOD_3_SR | 0.788 | 3.613014e-08 | Positive |

### Subject-specific global amplitude trends did not appear to be driven by core molecular clock amplitude or organization

We hypothesized that subject-specific differences in global amplitude could be explained by differences in the activity of the core molecular clock, considering both core clock amplitude, as well as organization (25).

First, we checked whether the subjects’ mean amplitude in a set of rhythmic core clock genes predicted their mean amplitude in the set of all other non-core clock rhythmic genes (**Figure 4A**). Some datasets (MUSCLE and BLOOD_2) showed a strong positive association between core clock and global amplitude, while one dataset (DERMIS_1) showed a nonsignificant inverse relationship, and others showed a modest or nonsignificant relationship. However, as we have found that there are shared global trends in amplitude across genes, we questioned if the relationship between global rhythm strength and core clock strength was stronger than the relationship we would expect between the amplitudes of a random collection of genes and global rhythm strength. **Figure 4B** shows that even in the datasets in which the core clock amplitude significantly predicted global amplitude, its predictive power did not exceed what would be expected from a size-matched set of randomly chosen rhythmic genes. In fact, in the skin datasets the core clock amplitude appeared to have substantially *less predictive* power than a random set of rhythmic genes of the same size.

**Figure 4:**
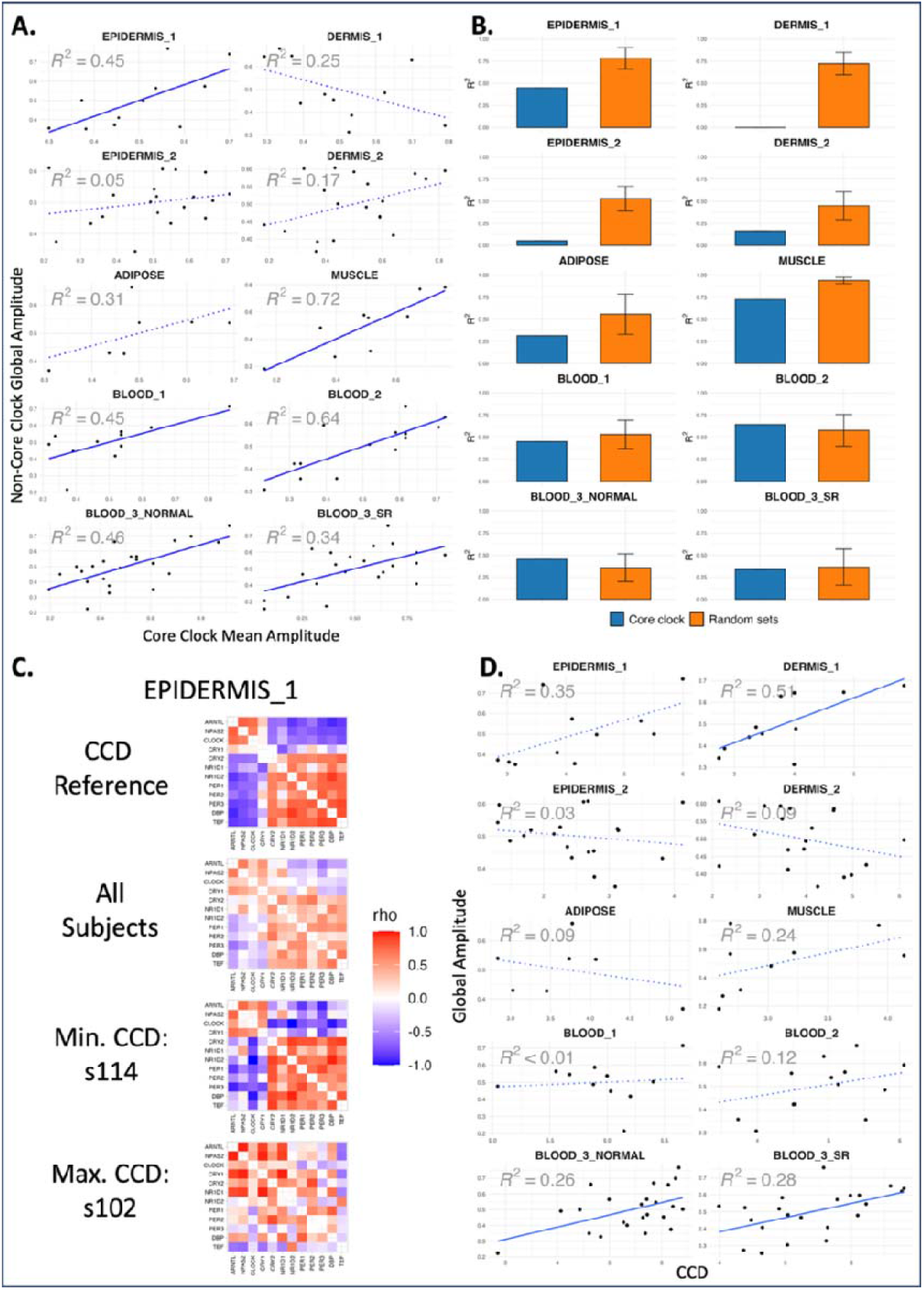
Comparison of metrics of core-clock amplitude and organization with global transcriptional amplitude measures. **A**. Subject-specific mean amplitude in rhythmic core molecular clock genes are compared to the mean amplitude in all other non-core-clock rhythmic genes. In some datasets, there is a significant correlation (shown with a solid blue line), while in others the correlation is non-significant (shown with a dotted blue line). **B**.) These plots show a comparison of predictive power (*R*^*2*^, recorded as 0 for negative correlations) in rhythmic core clock genes compared to a size-matched set of randomly selected set of genes in each dataset (with 100 runs total, to calculate standard error). Even in the datasets for which core clock mean amplitude predicted global amplitude in **Fig. 4A**, the predictive power did not substantially exceed what would be expected from a random set of rhythmic genes of the same size. **C**.) Example heatmaps from EPIDERMIS_1 of core clock correlations used for clock correlation distance (CCD): reference, across all subjects, the subject with the minimum CCD (closest to reference), the subject with the maximum CCD (further from reference). **D**.) Global amplitude compared to CCD for each subject (lower CCD = closer to reference). Solid blue lines show a significant correlation, while dotted blue lines show a non-significant correlation. We note that the only significant correlations were in the counterintuitive direction, with more *disorganized* core clocks predicting higher global amplitude.

Besides amplitude, we also considered the organization of the core clock. By “organization,” we mean whether or not the core molecular clock’s two transcriptional arms (activators, including CLOCK, ARNTL, NPAS2, and repressors, including PER and CRY family genes) show the expected phase-opposed, anti-correlated co-expression pattern across samples (activators peaking approximately 12h before repressors) (25). This type of core clock organization can be quantified using the clock correlation distance (CCD) measure described by Shilts et al. 2018 (26). The CCD is a reference-based distance that quantifies how closely the core-clock co-expression pattern in a given subject matches the canonical pattern (lower CCD = more closely matching, higher CCD = less closely matching, example in **Figure 4C**). We computed the CCD for each subject in each dataset using the *deltaccd* R package (27), and found that this measure of core clock organization generally did not predict global amplitude (**Figure 4D**). In fact, the only statistically significant associations (DERMIS_1, BLOOD_3_NORMAL, and BLOOD_3_SR) were in the counterintuitive direction, with more *disorganized* core clocks predicting higher global amplitude.

## Discussion

In this paper, we investigated whether or not some individuals have higher or lower amplitude in circadian gene expression *in general*, across the set of all rhythmic genes, and used permutation-based statistics to test this. The datasets we analyze all profiled purportedly healthy subjects with a relatively modest variation in age and all excluded shift workers. Despite this, we found that many subjects did indeed show a statistically significant global amplitude trend, and that this effect replicated across ten datasets from different tissues.

Still, our analysis has several potential limitations. The datasets we analyzed were limited in terms of the number of people in each cohort, with the largest group (BLOOD_3_NORMAL) having only 23 subjects, and the smallest group (ADIPOSE) having only 7 subjects. The datasets were also limited in the number of samples per individual, with the most densely sampled dataset (BLOOD_1) having 13-15 samples per subject, and the least densely sampled datasets (EPIDERMIS_2 / DERMIS_2) having only 4 samples per subject. However, although low sampling density undoubtedly limited our ability to estimate subject-specific rhythmic parameters for individual genes, this limitation should have been largely mitigated analytically, because all subjects within a given cohort were affected in a similar manner. It is also likely that, even though these studies generally excluded individuals with more “extreme” circadian phenotypes, subjects still differed in their circadian phase of entrainment. Importantly, once the set of cycling transcripts was defined for each study population, the acrophase of each transcript was allowed to vary for each subject when estimating that subject’s amplitude. Finally, although the original investigators appear to have followed standard experimental and preprocessing protocols before submitting their data to GEO, we cannot fully exclude the possibility that unknown experimental or technical artifacts influenced our analyses.

It is also important to note that each study focused on a relatively healthy cohort with a narrow age range. This is both a strength and a limitation. On one hand, our results demonstrate that even within an ostensibly homogeneous population, clear global differences in rhythm strength emerge between individuals. On the other hand, these differences may be modest compared to those that would arise in a more diverse population spanning a wider range of ages, sexes, and lifestyles (28–30). In such a population, the primary determinants of rhythm strength - and the pathways or factors with the greatest predictive value - might well differ

Additionally, the results we have presented were primarily statistical in nature. Although multiple plausible mechanisms could contribute—including lifestyle, endocrine signaling, body temperature, and inflammation (20,31–37)—we have not established a definitive or dominant mechanistic driver determining global-subject specific trends across tissues. We originally hypothesized that the expression of transcripts in the core molecular clock could be a driver of subject-specific differences in global amplitude, but the results of our analysis do not support this. However, while the amplitude or mean expression of this specific set of core circadian transcripts did not appear to explain global amplitude, it remains possible that genetic polymorphisms or epigenetic changes within the broad population do influence global amplitude. Moreover, it is possible that a broader cohort of subjects with more variation in core clock dynamics may have recovered a more marked effect. Indeed, it is likely that such effects are more significant in tumors or other disease states (11,26).

The core clock analysis results did suggest tissue specific differences as to the importance of core clock rhythms in determining global amplitude trends. In the blood datasets, the core clock was about equally as predictive of global amplitude as a randomly selected set of rhythmic genes, while in the skin datasets the core clock mean amplitude was substantially *less* predictive than a randomly selected set of rhythmic genes. Thus, while the global amplitude trend replicated across datasets from different tissues, the underlying biological controls and their relative importance may differ across tissues. Moreover, while the amplitude or mean value of this specific set of core circadian transcripts did not significantly predict global amplitude, it is possible that specific sequence polymorphisms or epigenetic modifications, rather than expression, do predict these changes. Furthermore, it is unclear how differences in global transcriptional amplitude relates to differences in other physiological rhythms, like those in body temperature or metabolism.

Besides the open question of underlying biological mechanism, there are related questions regarding the stability of the global amplitude trend throughout different parts of the body, over periods of time, and after perturbations or changes in behavior. We found a significant positive association between global amplitude in the epidermis and dermis of the same subjects, from skin samples taken at the same times, from the same skin sites, suggesting some persistence of global amplitude across different tissue subtypes that are close in proximity. Conversely, we did not find a significant association between global amplitude in blood samples from the same subjects, taken at different time points following either a week of sufficient sleep or a week of insufficient sleep, suggesting that in blood the global amplitude trend is either unstable over long periods of time, or unstable in response to changes in sleep habit, or both.

Despite these limitations, this work examines hypotheses that are widely invoked yet poorly established. We show that individuals exhibit clear, global, subject-specific tendencies in the strength of transcriptional rhythms. However, in younger and healthy cohorts, these differences are not primarily driven by variation in core clock rhythms. Understanding inter-individual variation in transcriptional amplitude will become increasingly important as efforts to personalize circadian medicine expand. By making our code publicly available, we hope to enable researchers to apply these tests to additional datasets and to investigate the biological mechanisms that underlie subject-specific differences in global amplitude.

